# Loss of *lrrk2* impairs dopamine catabolism, cell proliferation, and neuronal regeneration in the zebrafish brain

**DOI:** 10.1101/140608

**Authors:** Stefano Suzzi, Reiner Ahrendt, Stefan Hans, Svetlana A. Semenova, Saygın Bilican, Shady Sayed, Sylke Winkler, Sandra Spieß, Jan Kaslin, Pertti Panula, Michael Brand

## Abstract

*LRRK2* mutations are a major cause of Parkinson’s disease. Pathogenicity of LRRK2 loss-of-function is controversial, as knockout in rodents reportedly induces no brain-specific effects and knockdown studies in zebrafish are conflicting. Here we show that CRISPR/Cas9-engineered deletion of the ~60-kbp-long zebrafish *lrrk2* locus elicits a pleomorphic, albeit transient brain phenotype in maternal-zygotic mutants (mzLrrk2). Intriguingly, 11-month-old mzLrrk2 adults display increased dopamine and serotonin catabolism. Additionally, we find decreased mitosis in the larval brain and reduced stab injury-induced neuronal regeneration in the adult telencephalon. Finally, hypokinesia associates with loss of *lrrk2* in larvae. Our results demonstrate that *lrrk2* knockout has an early neurodevelopmental effect, and leads to perturbed dopamine and serotonin catabolism in a *LRRK2* knockout. We propose mzLrrk2 zebrafish as a valuable tool to study LRRK2 loss-of-function *in vivo*, and provide a link between LRRK2 and the control of basal cell proliferation in the brain that may become potentially critical upon challenges like brain injury.

## Introduction

The leucine-rich repeat kinase 2 (LRRK2/dardarin/PARK8) is a large multidomain protein and a bifunctional enzyme, displaying both kinase and GTPase activities [1]. *LRRK2* gene polymorphisms are the most recurrent genetic cause of familial and sporadic late-onset Parkinson’s disease (PD) reported so far, although the relative frequency of individual mutations varies with ethnicity [2]. *LRRK2* mutations cause pleomorphic and highly variable pathology, sometimes clinically indistinguishable from late-onset levodopa-sensitive idiopathic disease [3]. Typical features include selective dopaminergic cell loss and presence of Lewy bodies, abnormal protein aggregates associated with several neurodegenerative disorders [4].

*In vitro* evidence has initially suggested that pathogenic variants, including the relatively common G2019S substitution, confer toxicity via gain-of-function of the kinase domain [5, 6]. As a result, LRRK2 pharmacological inhibition attracts great interest in a therapeutic perspective. However, a loss-of-function contribution has not been ruled out. First, the current lack of reliable endogenous substrates or interaction partners makes it difficult to validate *in vitro* findings *in vivo*. More critically, mice overexpressing human wild-type [7] or mutant LRRK2 [8-13] do not generally recapitulate dopaminergic cell loss, unless transgene levels are artificially enhanced using strong promoters [11, 14-17]; Lewy body pathology has never been reported. Paradoxically, enhanced LRRK2 activity in G2019S knockin mice confers a hyperkinetic phenotype and seems protective against age-related motor impairment [18]. A further challenge comes from evidence that the G2385R variant, a risk factor for PD in the Chinese ethnicity [19, 20], reduces kinase activity [21, 22] and enhances LRRK2 degradation [21]. A similar dominant negative effect has been described also for the I2020T variant [23, 24], usually considered a gain-of-function. Finally, pathogenic variants may disrupt protein-protein interactions that may be essential in cell signaling pathways. Along this line, it has been shown that the substitutions R1441C/G/H, Y1699C, and I2020T, but not G2019S, reduce phosphorylation of LRRK2 residues S910/S935 in Swiss 3T3 cells, thus disrupting the interaction with 14-3-3 protein and causing non-14-3-3 bound LRRK2 to accumulate in inclusion body-like cytosolic pools [25]. Altogether, these observations back up an alternative loss-of-function hypothesis, bolstered by *Lrrk2* knockout in rodents being pathogenic in peripheral organs [26-28]. Remarkably, *Lrrk2* knockout mice develop PD-like pathology in the kidney, most prominently accumulation and aggregation of α-synuclein, but not in the brain [28]. Yet, LRRK2 deficiency in mice induces behavioral alterations similar to BAC human LRRK2 G2019S transgenic mice [29, 30].

These data emphasize the importance of additional knockout studies to elucidate LRRK2 function. *LRRK2* knockout models may help clarifying the molecular interactions that are disrupted by dysfunctional LRRK2, as well as potential side effects of LRRK2 inhibitors. The zebrafish is a valuable addition to rodent models due to its amenability to high-throughput studies and direct observation of disease mechanisms *in vivo*. In particular, the regenerative potential of the zebrafish adult brain [31] makes it a useful vertebrate system to determine which genetic programs might hinder neurodegeneration, with a view towards applying these insights in humans. Previous attempts to investigate zebrafish *lrrk2* gene function using morpholino oligonucleotides (MOs) yielded contradicting results. In one study, loss of diencephalic catecholaminergic (CA) neurons and locomotor defects in the larvae were described [32]. However, a subsequent study failed to reproduce the reported phenotype, even by using the same reagents and MOs [33]. Recently, a third paper rekindled the initial claims, describing a *lrrk2* MO-induced phenotype with strong developmental abnormalities [34]. These discrepancies revived concerns over the consistency of MO-induced knockdown to assess gene loss-of-function due to the variability and transiency of induced changes and the risk of off-target effects [35]. Although the analysis of MO-induced phenotypes may still provide useful information, their validation would inevitably entail the generation of reliable null alleles [36].

Using the clustered regularly interspaced short palindromic repeats (CRISPR)/CRISPR-associated protein-9 nuclease (Cas9) genome-editing tool, we report here the deletion of the ~60-kbp-long zebrafish *lrrk2* locus containing the entire open reading frame (ORF), resulting in an unambiguous null allele. To our knowledge, this is the first report describing LRRK2 deficiency in a vertebrate *in vivo* model where no residual or truncated protein is produced. We also report a second point mutant resulting in an early stop codon as a potential null allele. We characterized the phenotype of these alleles in the brain, as the organ of possibly highest relevance for PD. We find that maternal-zygotic *lrrk2* mutants display a pleomorphic, but transient neurodevelopmental phenotype, including increased apoptosis, delayed myelination, reduced and morphologically abnormal microglia/leukocytes, and reduced catecholaminergic neurons. We also find a correlation between hypokinesia and loss of *lrrk2* in larvae. Importantly, we also find perturbed amine catabolism in progressively older adult animals. Finally, we observe decreased mitosis in the larval brain and impaired neuronal regeneration after stabbing the adult telencephalon. Our results suggest a link between zebrafish Lrrk2 and the control of cell proliferation in the brain, with crucial implications for the self-healing capacity upon lesion.

## Results

### Deletion of the entire lrrk2 locus using CRISPR/Cas9

Syntenic analysis using the assemblies of the human and zebrafish genomes (GRCh38p.7 and GRCz10, respectively) revealed the conservation of *SLC2A13* as the downstream neighbor of *LRRK2* in both species. Moreover, duplication of the zebrafish *lrrk2* locus is not reported. The human LRRK2 and zebrafish Lrrk2 proteins share the same domain structure, with the kinase domain displaying the highest degree of conservation (Figure 1a); three of the four sites of pathogenic substitutions in humans are fully conserved (Table 1). Together, these data suggest that human *LRRK2* and zebrafish *lrrk2* are truly orthologous genes.

**Figure 1.**
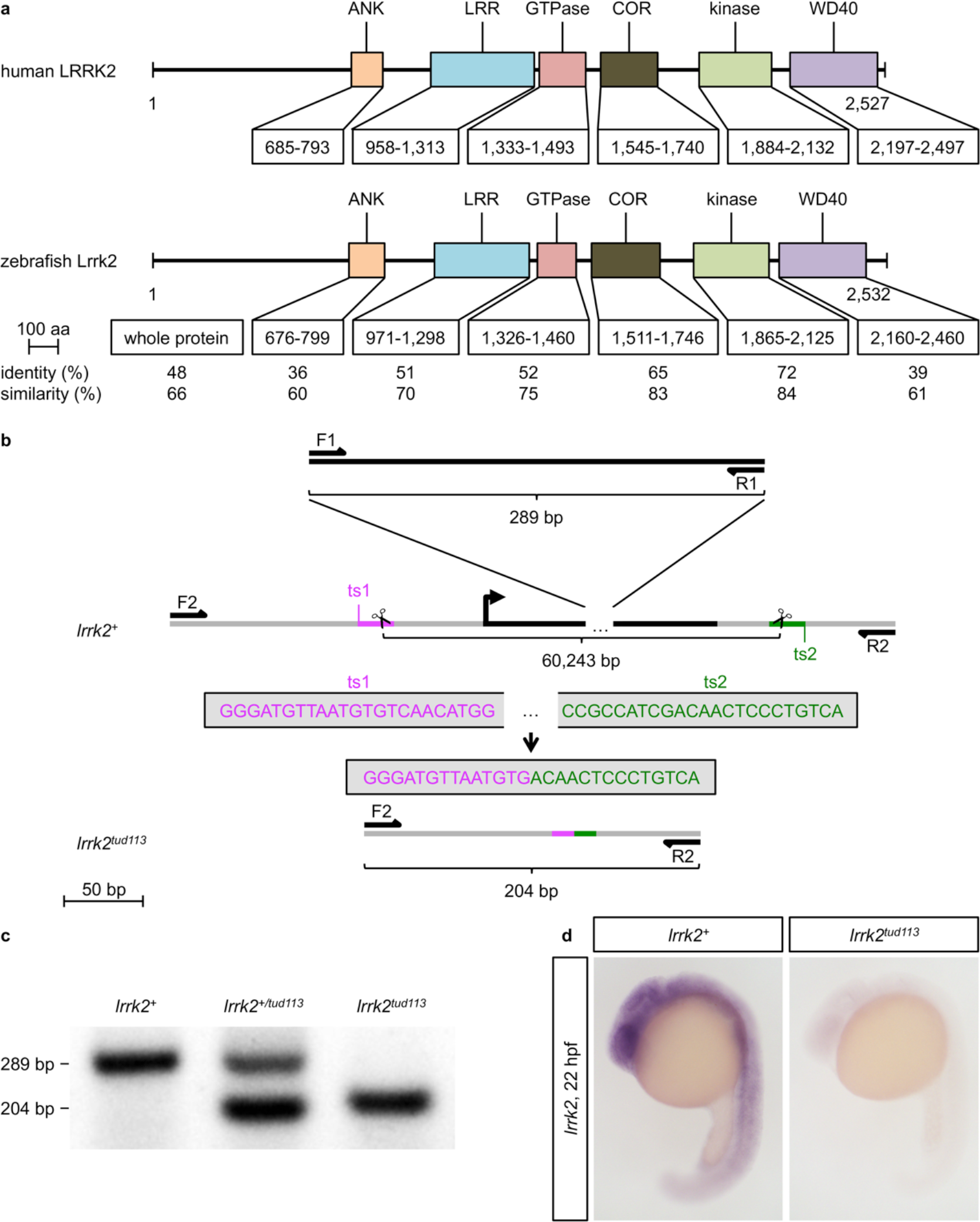
Homology between the human LRRK2 and the zebrafish Lrrk2 proteins and generation of the zebrafish *lrrk2^tud113^
* allele. (a) Alignment analysis of the whole sequence and the individual domains of human LRRK2 (NP_940980) and zebrafish Lrrk2 (NP_001188385) proteins reveals a high degree of conservation of the catalytic core. The percentages of identity (same residues at the same positions in the alignment) and similarity (identical residues plus conservative substitutions) are indicated. Abbreviations: ANK, ankyrin domain; COR, C-terminal of Ras of complex proteins; LRR, leucine-rich repeat domain. (b) Scheme reproducing the targeting and screening strategy. The *lrrk2* open reading frame (ORF) is highlighted in black; F1, R1: *lrrk2 ORF*-specific primers; F2, R2*: lrrk2 ORF*-flanking primers; ts1 (magenta), ts2 (green): gRNA target sites (ts). (c) gap-PCR analysis of genomic DNA from wild-type (*lrrk2^+^
*), heterozygous (*lrrk2^+/tud113^
*), and homozygous mutant (*lrrk2^tud113^
*) individuals. F1 and R1 amplify a 289-bp-long product, F2 and R2 a 204-bp-long product. (d) *lrrk2 in situ* hybridization confirming the complete absence of *lrrk2* expression in 22-hpf *lrrk2^tud113^
* embryos.

**Table 1.** Sites of pathogenic amino acid substitutions in human LRRK2 and corresponding residues in zebrafish Lrrk2.

| substitution in humans | corresponding site in zebrafish | protein domain |
| --- | --- | --- |
| p.Arg1441Cys/Gly/His |  |  |
| p.Tyr1699Cys | Tyr1685 | COR |
| p.Gly2019Ser | Gly2009 | kinase |
| p.Ile2020Thr | Ile2010 | kinase |

Data retrieved from the Parkinson Disease Mutation Database (PDmutDB, http://www.molgen.vib-ua.be/PDmutDB) [37, 38].

To study the gene function of zebrafish *lrrk2 in vivo* and potentially resolve the controversy arising from previous MO studies [32-34], we removed the *lrrk2* locus containing the entire ORF. This approach has two main advantages. First, potential side effects of frameshift mutations, including cellular stress due to aberrant transcripts or truncated protein products with residual or new function, are excluded. Second, identification of mutation carriers is straightforward. To achieve full deletion of the *lrrk2* locus, two CRISPR/Cas9 target sites flanking the 60 kb-long ORF were chosen (Figure 1b). To identify deletion alleles, a gap-PCR strategy was devised, with primers amplifying a 289-bp-long amplicon inside the target region duplexed with flanking primers, unable to direct amplification unless a deletion brings them in sufficient reciprocal proximity (Figure 1b, c). The selected founder produced offspring where the flanking primers amplified a 204-bp-long product, revealing a 60,243-bp-long targeted deletion (c.−61_*42del according to the Human Genome Variation Society guidelines [39]; henceforth this allele is referred to as “tud113”) as confirmed by sequencing. The complete absence of *lrrk2* expression in homozygous tud113 mutants was verified via *in situ* hybridization (Figure 1d).

F1 heterozygous fish were incrossed to obtain F2 homozygous tud113 mutants and wild types. To exclude any effect from the existing maternal wild-type *lrrk2* transcripts (Figure S1a), F2 fish were further incrossed to obtain homozygous mutants that developed from homozygous mothers (maternal-zygotic homozygous tud113 mutants, henceforth referred to as “mzLrrk2”) and wild-type (wt) control lines. In striking contrast with published MO-induced phenotypes [32, 34], mzLrrk2 individuals develop normally, are viable, and reach sexual maturity at the same age as controls, with both females and males being fertile. During zebrafish development, *lrrk2* expression is ubiquitous until 24 hours post-fertilization (hpf), then gradually restricts to the head (Figure S1); *lrrk2* transcripts are relatively scant in whole-body 5- and 10-dpf larvae, as measured by RT-qPCR (not shown). In the adult brain, *lrrk2* expression is present throughout the whole organ (Figure S2). The time course of *lrrk2* expression in the zebrafish brain thus closely mirrors the situation in rodents, where *Lrrk2* is broadly expressed in the embryonic, postnatal, and adult brain, though at low levels compared to other peripheral tissues [40-42]. To exclude possible compensation from the paralogous *lrrk1* gene, we also characterized *lrrk1* expression in developing embryos (Figure S3a–f’). Compared to *lrrk2* (Figure S1a–i), *lrrk1* expression is markedly tissue-restricted. Moreover, *lrrk1* is not upregulated in mzLrrk2 larvae either at 5 or at 10 days post-fertilization (dpf) as determined by RT-qPCR (Figure S3g).

### Pleomorphic but transient brain phenotype in mzLrrk2 fish

Because the link between *LRRK2* and PD in humans implies a critical role in brain function, we characterized mzLrrk2 fish with regard to the brain phenotype at both larval (5 and 10 dpf) and adult (6 and 11 months, mo) stages. Using TUNEL assay, we measured a threefold increase of the cell death rate in mzLrrk2 brains at 5 dpf (anterior: *P*=0.0061; middle: *P*<0.0001; Figure 2a, a’), but not at 10 dpf (Figure 2b, b’). Unlike previous studies [32, 34], no clear sign of neuronal loss was evident after HuC/D staining at either 5 or 10 dpf (Figure 2c, d). Likewise, the commissural acetylated Tubulin+ axons in the tectum appeared normal at both larval time points (Figure 2e, f). To further investigate the consequences of Lrrk2 deficiency on neural development, Claudin k staining was performed to visualize myelination. Claudin k expression was delayed in the ventromedial hindbrain of mzLrrk2 at 5 dpf (Figure 2g), but normal at 10 dpf (Figure 2h). In summary, our data show that loss of *lrrk2* causes early, but transient defects, including increased apoptosis and delayed myelination, albeit no obvious signs of neurodegeneration.

**Figure 2.**
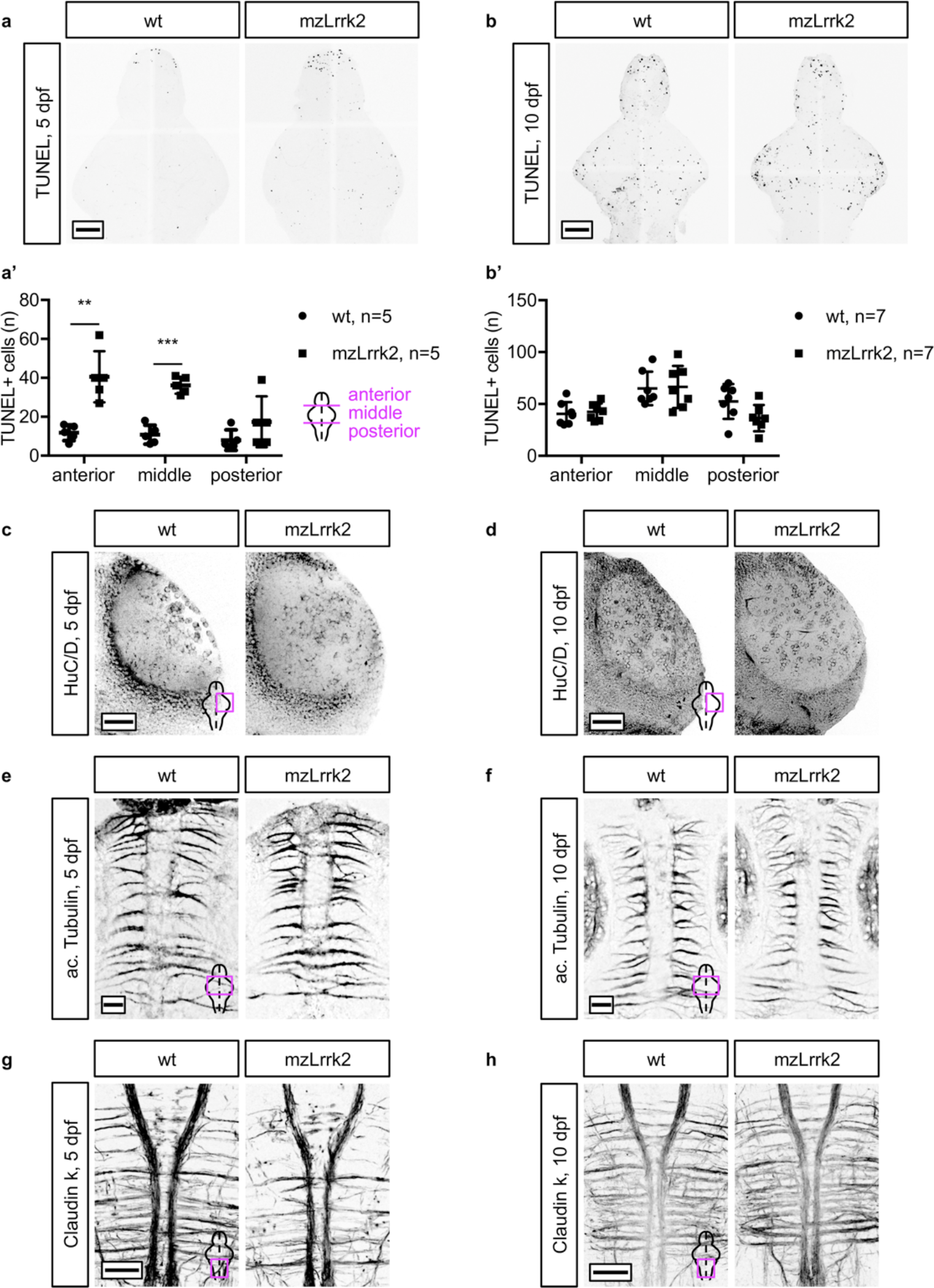
Initial neurodevelopmental abnormalities are subsequently compensated. (a, b) TUNEL assay to visualize cell death in the brain. Quantification was carried out over the whole brain, subdivided into anterior (telencephalon), middle (diencephalon, mesencephalon), and posterior (rhombencephalon) portions. The number of apoptotic cells in mzLrrk2 brains is increased at 5 dpf (a, a’), but matched wt levels at 10 dpf (b, b’). (c, d) HuC/D immunohistochemistry to label mature neurons and (e, f) acetylated Tubulin immunohistochemistry to mark the axonal network at 5 (c, e) and 10 dpf (d, f) does not reveal overt signs of degeneration in the optic tectum. (g, h) Claudin k immunohistochemistry shows a delay in myelination in the ventral hindbrain at 5 dpf (g) that is later compensated at 10 dpf (h). (a’, b’) Plots represent means±s.d. Statistical analyses: (a’, b’) two-tailed Student’s *t*-test. Scale bars: (a, b, g, h) 100 *µ*m; (c, d) 50 *µ*m; (e, f) 20 µm.

Neuroinflammation is a concurring factor in many neurodegenerative conditions, including PD [43]. Increasing evidence implicates *LRRK2* in microglia function [44]. Because the shape of microglia may be indicative of their activation state, we used the pan-leukocyte marker L-Plastin and examined the brain microglia/leukocyte number and morphology (Figure S4a). The overall number was reduced by about one third in 5-dpf mzLrrk2 larvae (Figure 3a, a’; *P*=0.0273). The trend was maintained until 10 dpf, though without reaching statistical significance (*P*=0.0797; Figure 3b, b’). For each brain, average microglia/leukocyte volume, surface, ramification, average branch length, maximum branch length, and longest shortest path were used to study the variance across samples via principal component analysis. While little separation existed between the 5- and 10-dpf wt samples, the 5- and 10-dpf mzLrrk2 samples segregated, with 10-dpf mzLrrk2 clustering with controls (Figure 3c). In particular, mzLrrk2 microglia/leukocytes were on average larger and more ramified at 5 dpf, to then become smaller and less complex, comparable to the wt, at 10 dpf (Figure S4b– o). To test whether even mild morphological alterations correlated with impaired leukocyte function, 10-dpf larvae were treated with 12-*O*-tetradecanoylphorbol-13-acetate (TPA) to induce systemic acute inflammation [45] for 2 h prior to killing. Although treatment was not sufficient to elicit a substantial effect on the brain (not shown), significantly quenched leukocytosis was found in the tail (*P*=0.0368; Figure 3d, d’). In conclusion, our data suggest that zebrafish Lrrk2 plays a transient role in leukocyte biology, including a response to proinflammatory stimuli, and additionally hint that the recreation of pathological conditions in animal models, such as inflammation, may be an essential expedient for a thorough understanding of LRRK2 function.

**Figure 3.**
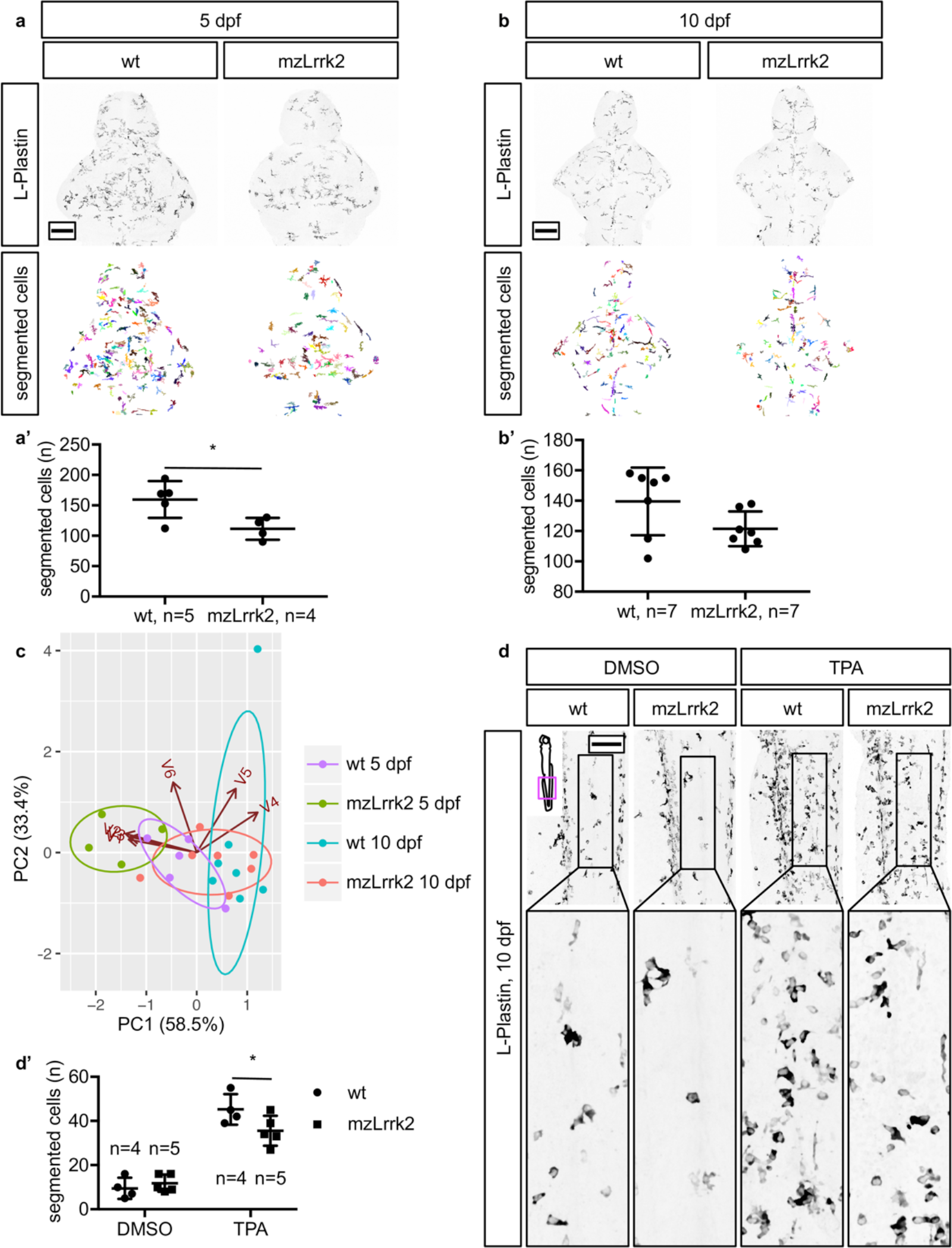
Microglia/leukocyte cell number and morphology are altered in mzLrrk2 mutants. (a, a’, b, b’) Quantification of segmented L-Plastin+ cell number in the brain. The number of microglia/leukocytes was reduced in mzLrrk2 brains at 5 dpf (a, a’), but comparable to controls at 10 dpf (b, b’). (c) Principal component analysis of brain microglia/leukocyte morphology. For each animal, the average morphological parameters were considered; standardized data were used. The samples (dots) and the original variables (arrows) are projected on the 2D plane defined by PC1 and PC2: V1, volume; V2, surface; V3, ramification; V4, average branch length; V5, maximum branch length; V6, largest shortest path. PC1 opposes cell size (V1, V2) and ramification (V3) to branch extension (V4, V5); PC2 represents overall cell complexity (V4, V5, and V6). Within brackets, the percentage of total variance explained. Note the poor separation between mzLrrk2 and wt samples at 10 dpf. (d, d’) Reduced leukocyte response in the tail of 10-dpf mzLrrk2 larvae after 2 h exposure to 12-*O*-tetradecanoylphorbol-13-acetate (TPA) to induce systemic acute inflammation. Segmented objects were quantified within a constant 566.79×124.54 µm area caudal to the anus and comprised between the dorsal longitudinal anastomotic vessel and the caudal artery. Sample n are indicated in the graph (d’). (a’, b’, d’) Plots represent means±s.d. Statistical analyses: (a’, b’) two-tailed Student’s *t*-test; (d’) one-tailed Student’s *t*-test. (a, b, d) Scale bars: 100 µm.

### Intact CA system but perturbed amine catabolism in older fish

Because catecholaminergic (CA) neurons are the most clinically relevant target of PD, special attention was paid to the CA system. CA cell populations along the rostro-caudal axis from the olfactory bulb to the *locus cœruleus* were visualized via tyrosine hydroxylase (TH) immunohistochemistry and denominated after the nomenclature introduced previously [46] (Figure 4a). Zebrafish possess two paralogous *th* genes: *th1* and *th2* [47]. Because commercially available anti-TH antibodies only recognize TH1, but not TH2 protein [48], we analyzed the zebrafish CA system by double staining with an anti-TH1 antibody and a recently characterized pan-TH antibody to also identify TH2+ cells by exclusion (Figure 4a’). Additionally, the levels of biogenic amines and their catabolites (Figure 4b) were measured via electrochemical detection coupled with high performance liquid chromatography. We observed that no TH+ cell population was missing or overtly altered in mzLrrk2 fish. However, at 5 dpf, mzLrrk2 brains displayed lower numbers of TH+ cells in discrete populations: olfactory bulb (pop. 1, *P*=0.0313); telencephalic complex (pop. 2, *P*=0.0322); diencephalic complex (pop. 5, 6, 11, *P*=0.0042); and paraventricular organ, *partes intermedia* and *posterior* (pop. 9, 10 *P*=0.0042; Figure 4c, c’). The net effect was a 20% reduction of the overall number of TH+ cells (not shown). Because mzLrrk2 brains showed a higher cell death rate at 5 dpf (Figure 2a, a’), TUNEL assay was combined with TH immunohistochemistry to investigate whether CA neurons were selectively affected. However, virtually no colocalization was found (not shown). Consistently, the amine catabolism appeared normal (Figure 4d). At 10 dpf, only the hypothalamic complex (pop. 13, *P*=0.0403) and, mildly, the paraventricular organ, *pars posterior* (pop. 10b, *P*=0.0496) were affected; in contrast, an increase in the preoptic area, *pars posterior* (pop. 4, *P*=0.0138) was measured (Figure 4e, e’). In general, the overall number of TH+ cells did not differ from the controls (not shown). Intriguingly, although the dopamine catabolism was unaffected, a higher concentration of 5-hydroxyindoleacetic acid, a catabolite of serotonin, was measured (*P*=0.0190; Figure 4f). Because brain-specific effects could be masked in whole-larvae homogenates, the activity of monoamine oxidase (MAO), one of the two major catabolizing enzymes, was histochemically visualized in 5- and 10-dpf brains (Figure 4g, h). However, no difference was observed between mzLrrk2 and controls.

**Figure 4.**
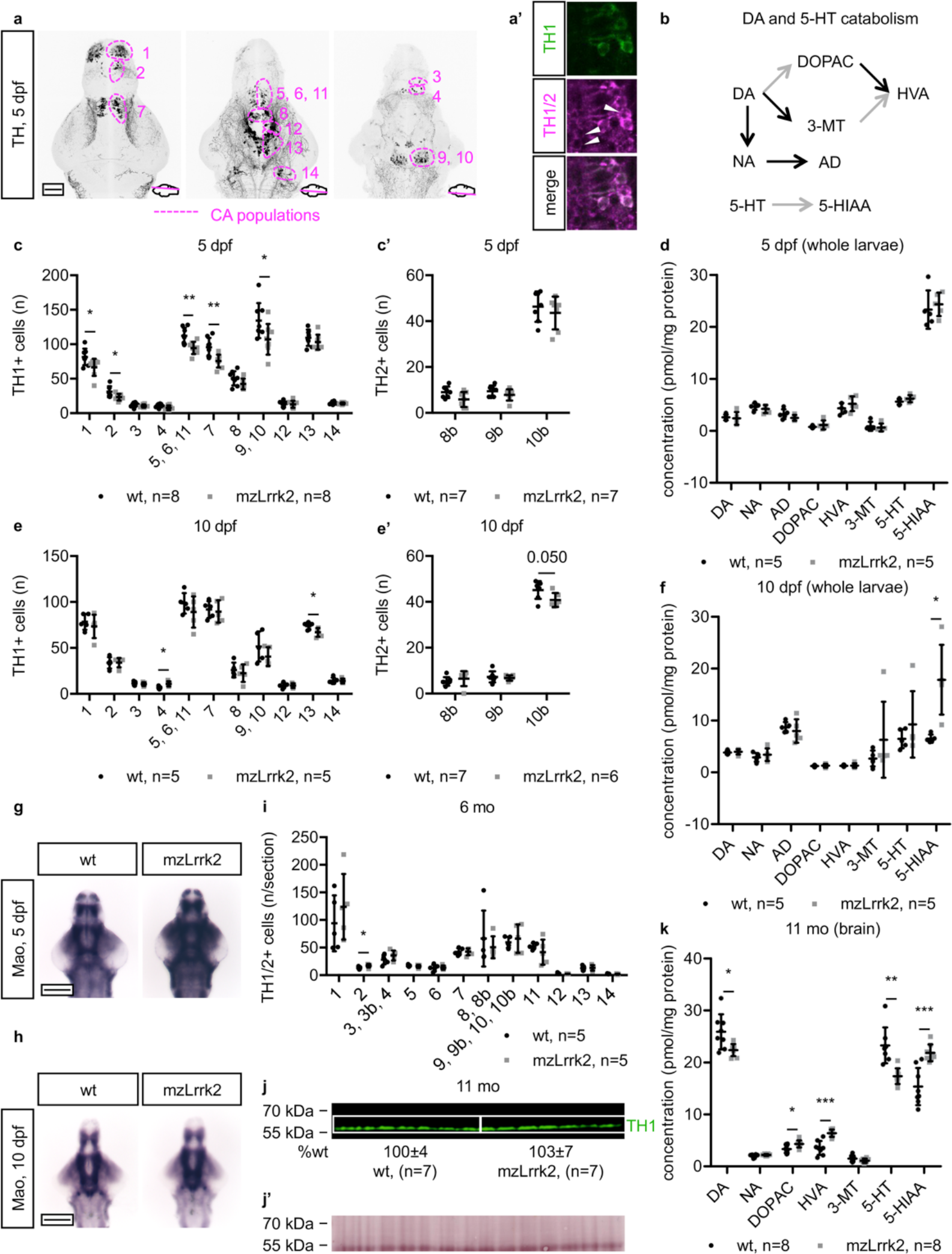
Loss of *lrrk2* causes transient developmental defects of the catecholaminergic system and perturbed amine catabolism in older fish. (a) The zebrafish catecholaminergic (CA) cell populations in the rostro-caudal axis from the olfactory bulb to the *locus cœruleus* as revealed by tyrosine hydroxylase 1 (TH1) immunohistochemistry at 5 dpf. Pop. 1: olfactory bulb; pop. 2: telencephalic complex; pop. 3: preoptic area, *pars anterior*; pop. 4: preoptic area, *pars posterior*; pop. 5, 6, 11: diencephalic complex; pop. 7: pretectal area; pop. 8: paraventricular organ, *pars anterior*; pop. 9: paraventricular organ, *pars intermedia*; pop. 10: paraventricular organ, *pars posterior*; pop. 12: posterior tuberal nucleus/posterior tuberculum; pop. 13: hypothalamic complex; pop. 14: *locus cœruleus*. (a’) Combination of the anti-TH1 antibody with the pan-TH antibody allows the identification of TH2+ cells (white arrowheads) by exclusion. TH2+ cells are found within the TH1 pop. 8, 9, and 10 in the paraventricular organ, thereby constituting the TH2 pop. 8b, 9b, 10b. (b) Simplified scheme of the catabolism of dopamine and serotonin. Each arrow represents a distinct enzymatically-catalyzed step. Grey arrows indicate reactions catalyzed by the combined action of monoamine oxidase/aldehyde dehydrogenase. Abbreviations: 3-MT, 3-methoxytyramine; 5-HIAA, 5-hydroxyindoleacetic acid; 5-HT, serotonin; AD, adrenalin; DA, dopamine; DOPA, 3,4-dihydroxyphenylacetic acid; HVA, homovanillic acid; NA, noradrenalin. (c–h) Quantification of TH1+ (c, e) and TH2+ cells (c’, e’) in the brain, quantification of amine catabolites in whole larvae (d, f), and histochemical visualization of monoamine oxidase activity in the brain (g, h) at 5 dpf (c–d, g) and 10 dpf (e–f, h). Early defects in discrete CA cell populations at 5 dpf are resolved by 10 dpf. The dopamine catabolism is normal at both time points. (i) Quantification of TH1/2+ cells in 6-mo brains and (j) quantification of TH1 protein levels in 11-mo brains show intact CA system in adult fish. (j’) Total protein stain as loading control for the immunoblot in (j). (k) Analysis of amine catabolism in 11-mo brains reveals increased dopamine and serotonin degradation. (c–f, i, k) Plots represent means±s.d. (j) Protein levels are reported as means±s.d. Statistical analyses: (c pop. 2–14, c’, d DA–HVA, d 5-HT, d 5-HIAA, e pop. 1, e 3–14, e’, f DA–HVA, f 5-HIAA, i pop.1, i pop. 3–11, i pop. 13, i pop. 14, j, k DA, k DOPAC–5-HIAA) two-tailed Student’s *t*-test; (c pop. 1, d 3-MT, e pop. 2, f 5-HT, i pop. 2, i pop. 12) two-tailed Mann-Whitney’s *U*-test. (a, g, h) Scale bars: 100 *µ*m.

In the adult brain, the CA system appeared structurally intact at 6 mo, with a modest increase in cell number in the telencephalic complex (pop. 2, *P*=0.0472; Figure 4i). Furthermore, TH1 protein levels at 11 mo were normal (Figure 4j). However, the amine catabolism in the brain was clearly perturbed in the older fish (Figure 4k). Specifically, a significant decrease of both dopamine (*P*=0.0216) and serotonin (*P*=0.0013) was found and, consistently, a significant increase in their catabolites 3,4-dihydroxyphenylacetic acid (*P*=0.0187), homovanillic acid (*P*=0.0001), and 5-hydroxyindoleacetic acid (*P*=0.0004), all products of MAO activity, was detected. However, MAO protein levels appeared unaltered, as determined on both gene expressional (Figure S5a) and biochemical level (Figure S5b). Nonetheless, the levels of the dopamine catabolite 3-methoxytyramine, product of catechol-*O*-methyltransferase activity, were not significantly different (*P*=0.1250; Figure 4k). Because monoamine oxidase and catechol-*O*-methyltransferase are the major enzymes responsible for catecholamine catabolism in the brain, we conclude that the neurochemical signatures observed were ascribable to MAO activity. In conclusion, although the cellular composition of the CA system appears morphologically unaltered, 11-mo mzLrrk2 fish show increased MAO-mediated amine degradation in the brain.

### Decreased mitosis rate in the larval brain

Several lines of evidence point towards an involvement of mammalian LRRK2 in cell proliferation and differentiation [49]. Impaired neurogenesis and neurite outgrowth were reported in BAC human LRRK2 G2019S transgenic mice [50]. Knockout studies have, however, led to ambiguous findings: in mice, DCX+ neuroblasts in the dentate gyrus were increased in one study [51], but unaffected in another [29]. To assess cell proliferation in the zebrafish mzLrrk2 brain, we used phospho-histone H3 (pH3) as a marker for mitotic cells. At 5 dpf, mzLrrk2 brains displayed a significantly reduced number of mitotic cells in the anterior brain only (Figure 5a, a’; *P*=0.0052). However, the effect at 10 dpf accentuated, consisting in a 50% decrease in the anterior (*P*=0.0025) and middle brain portions (*P*=0.0062; Figure 5b, b’). The phenotype is clearly apparent only in maternal-zygotic individuals (Figure S6). Intriguingly, the number of cells in S phase of the cell cycle did not differ between mzLrrk2 and controls (anterior: *P*=0.6453; middle: *P*=0.1455; posterior: *P*=0.3613; Figure 5c), suggesting defects either in the progression or in the regulation of the cell cycle length. This is in line with previous evidence showing impaired cell cycle progression, survival and differentiation of human mesencephalic neural progenitor cells upon *LRRK2* knockdown [52]. Altered mitotic rate is not sufficient to cause overall altered brain size or morphology (not shown). Remarkably, the phenotype is merely emerging at 5 dpf and exacerbates by 10 dpf, i.e. during a time window when neural development slows down considerably, judging from the about nine-fold drop in the absolute number of total pH3+ cells in wt brains (5 dpf: 281.3±26.7; 10 dpf: 33.9±8.7).

**Figure 5.**
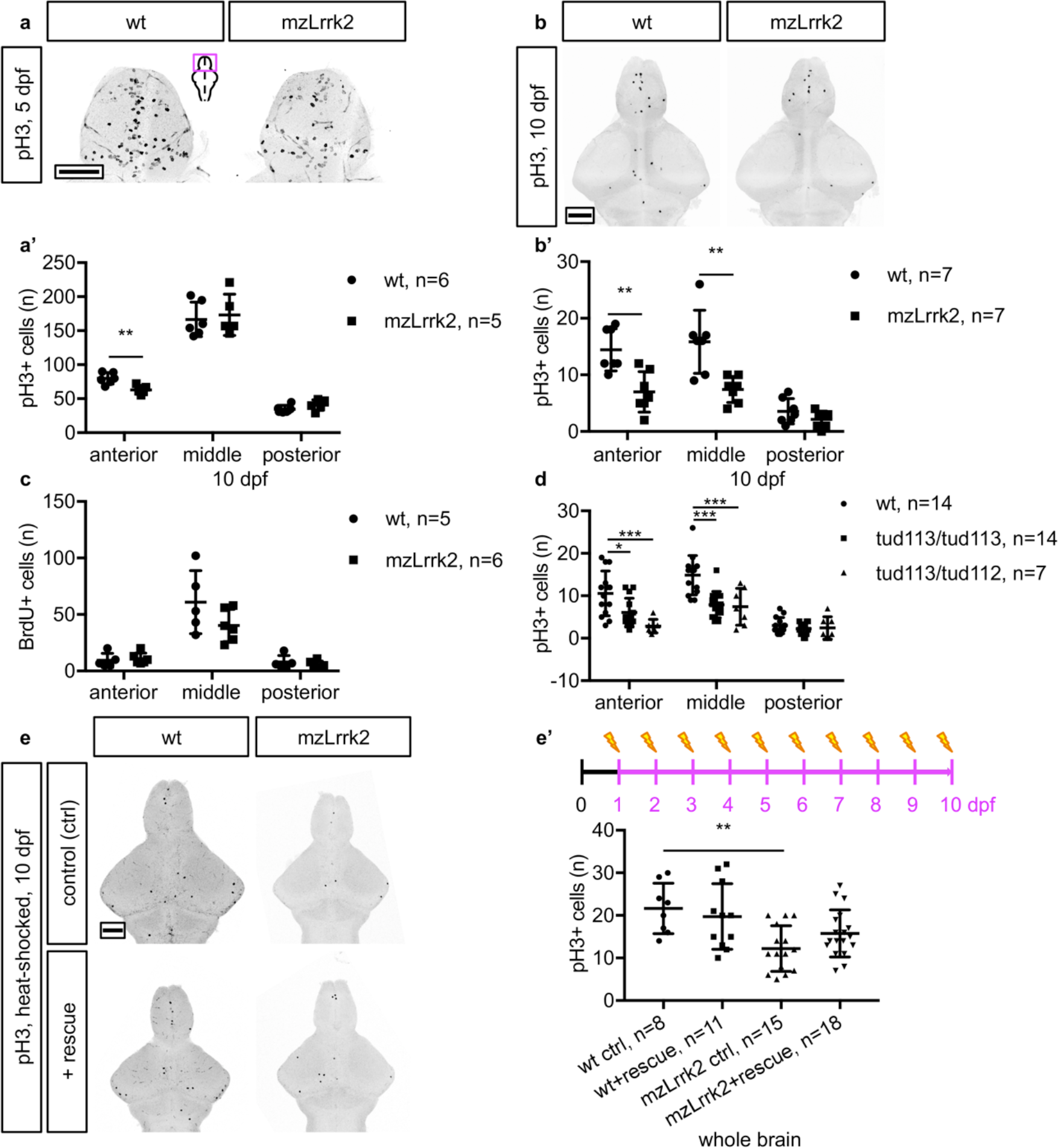
Loss of *lrrk2* results in reduced mitosis in the larval brain. (a, b) Phospho-histone H3 (pH3) immunohistochemistry to label mitotic cells. The mitosis rate is slightly reduced in the anterior portion only at 5dpf (a, a’), halved over the whole organ at 10 dpf (b, b’). (c) BrdU immunohistochemistry to label cells in the S phase of the cell cycle. To this aim, a 30 min BrdU pulse was delivered prior to killing. No significant difference between mzLrrk2 and controls was apparent. (d) Combination of the *lrrk2^tud113^
* and *lrrk2^tud112^
* alleles recapitulates the hypoproliferative phenotype in 10-dpf maternal-zygotic tud113 brains. (e, e’) Conditional expression of a Lrrk2 fragment (wt+rescue, mzLrrk2+rescue) containing the catalytic core rescues the hypoproliferative phenotype in 10-dpf mzLrrk2 brains (mzLrrk2+rescue versus mzLrrk2 controls, ctrl). (a’, b’, c, d, e’) Plots represent means±s.d. Statistical analyses: (a’, b’, c anterior, c middle) two-tailed Student’s *t*-test; (c posterior) two-tailed Mann-Whitney’s *U*-test; (d) one-way ANOVA followed by Tukey’s *post hoc* test; (e’) one-way ANOVA followed by Dunn-Šidák’s correction for multiple comparisons. Multiple comparisons: (e’) wt ctrl vs. wt+rescue; wt ctrl vs. mzLrrk2 ctrl; wt ctrl vs. mzLrrk2+rescue; mzLrrk2 ctrl vs. mzLrrk2+rescue. Scale bars: (a, b, e) 100 *µ*m.

To further validate our findings we carried out a complementation assay by combining the *lrrk2^tud113^
* allele with another *lrrk2* null allele to test recapitulation of the homozygous tud113 phenotype. We used a *N*-ethyl-*N*-nitrosourea-mutagenized *lrrk2* line (henceforth referred to as “tud112”) generated in our lab [53] that carries a frameshift mutation (c.3972+2T>C) causing a premature stop within the LRR domain-coding region (p.(Ile1252AlafsTer9); Figure S7). Maternal-zygotic tud112 mutants demonstrated a similar decrease of mitosis in 8-dpf brains (Figure S7d). Upon crossing homozygous tud113 females with tud112 males, we confirmed the hypoproliferative phenotype also in 10 dpf-transheterozygotes (compared to wt, anterior: *P*=0.0002; middle: *P*=0.0007; Figure 5d).

Furthermore, we sought to ‘rescue’ Lrrk2 function by overexpressing a Lrrk2 protein fragment containing the catalytic core in mzLrrk2 mutants. To this aim, we generated a Tol2 transgenic line expressing a *lrrk2* rescue construct (c.3009_7130) along with an mCherry reporter under the heat-inducible *hsp70l* promoter (*Tg(hsp70l:mCherry-T2A-lrrk2(c.3009_7130)-Myc)^tud114^
*, henceforth referred to as “rescue”; Figure S8). The rescue line was combined with the tud113 line to determine if the maternal-zygotic phenotype can be rescued. Reconstituted fish were heat-shocked daily for 4 h from 1 to 10 dpf and analyzed for mitosis in the brain. Compared to wt controls, the overall number of pH3+ cells in the mzLrrk2+rescue recovered (*P*=0.1051), as opposed to mzLrrk2 controls (*P*=0.0034; Figure 5e, e’). This finding indicates that overexpression of a functional Lrrk2 protein fragment can rescue the mzLrrk2 phenotype. Altogether, our data reveal an implication of Lrrk2 in the control of cell proliferation in the developing brain.

### Impaired neuronal regeneration in the adult telencephalon

Zebrafish have a remarkable capacity to regenerate lost or damaged tissues, including the telencephalon [54] and cerebellum [55]. Since regeneration recapitulates several aspects of embryonic development, we wondered whether loss of *lrrk2*, responsible for decreased cell proliferation in the larval brain, also affects reactive neurogenesis and regeneration in the adult brain. In unlesioned 6-mo old mzLrrk2 brains, we observed that proliferating cells, stained for the proliferation marker PCNA, are mildly reduced in the dorsal telencephalic niche, and significantly so in the dorso-posterior (Dp) area (*P*=0.0272; Figure S9). To address reactive proliferation and neurogenesis, 6-mo fish were injured using a stab lesion paradigm as previously described [54] (Figure 6a) and analyzed using a BrdU pulse-chase assay. Reactive proliferation at 3 days post-lesion (dpl) did not differ between mzLrrk2 and controls (Figure 6b). However, mzLrrk2 brains displayed on average 30% less HuC/D+/BrdU+ neurons at 21 dpl (*P*=0.0262; Figure 6c, c’). In contrast, neurogenesis in the unlesioned hemisphere was normal (*P*=0.284). Taken together, our results suggest a role for Lrrk2 in neuronal regeneration of the lesioned adult telencephalon.

**Figure 6.**
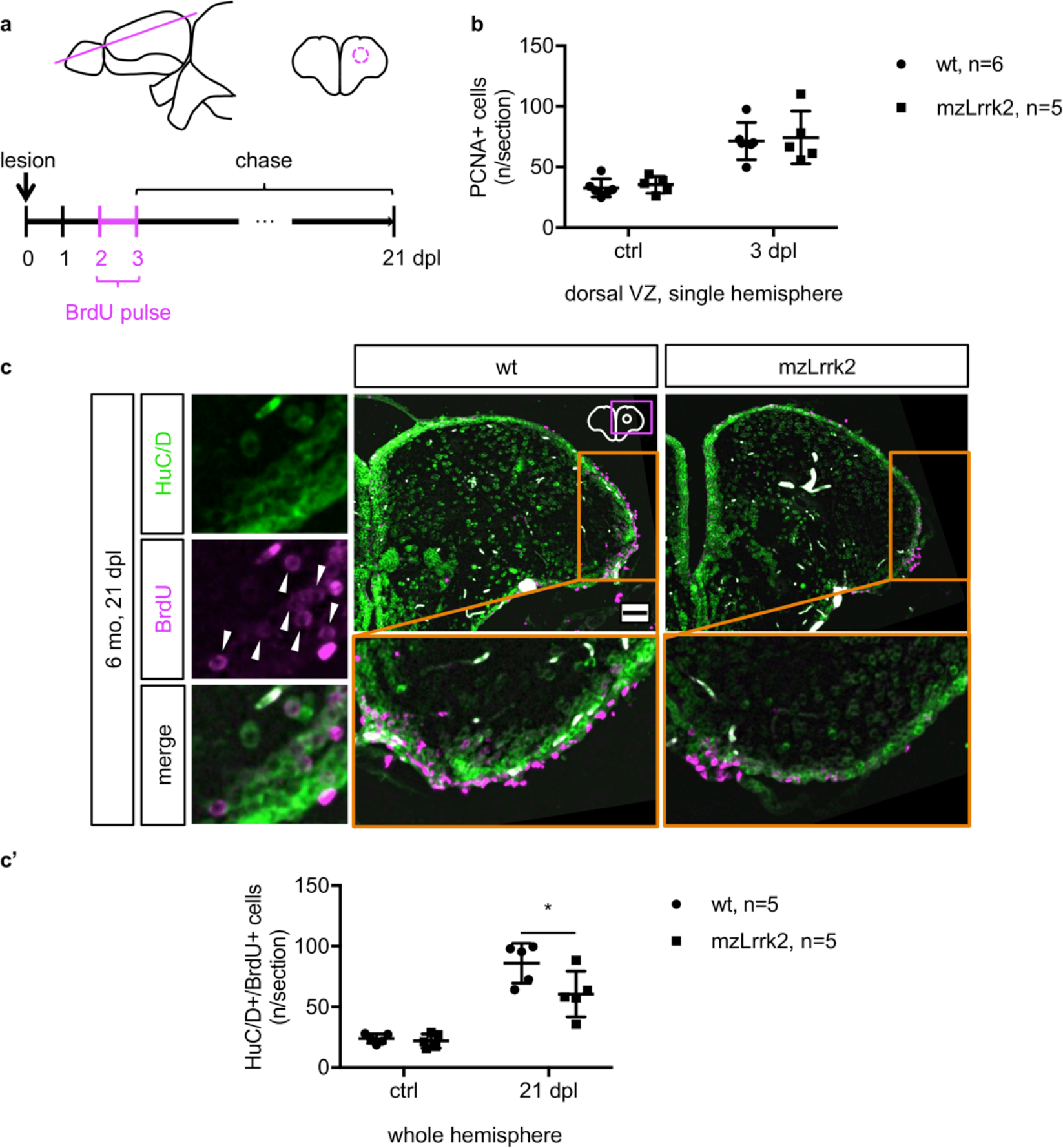
Loss of *lrrk2* impairs neuronal regeneration upon stab injury of adult telencephalon. (a) A unilateral stab injury was inflicted to the 6-mo adult telencephalon. To identify newborn neurons, a BrdU pulse was delivered from 2 to 3 days post-lesion (dpl) and incorporating cells analyzed at 21 dpl. (b) PCNA immunohistochemistry to examine reactive proliferation in radial glia stem cells of the ventricular zone (VZ) at 3 dpl. The lesioned hemisphere (3 dpl) was compared to the unlesioned hemisphere as control (ctrl) but no difference was observed (c, c’). HuC/D/BrdU double labeling to assess neurogenesis at 21 dpl. Double-positive cells are indicated by white arrowheads in the insets. Quantification was carried out through the entire parenchyma of the lesioned (21 dpl) and ctrl hemispheres. Neurogenesis is reduced in the mzLrrk2 brains. Orange-boxed insets are rotated by 90°. (b, c’) Plots represent means±s.d. Statistical analyses: one-tailed Student’s *t*-test. (c) Scale bar: 50 µm.

### Association between loss of lrrk2 and hypokinesia in larvae

PD patients are characterized by a motor syndrome comprising resting tremor, rigidity, postural instability, and bradykinesia, or slowness of movement. To investigate motor ability after loss of *lrrk2*, spontaneous swimming activity of 5- and 10-dpf larvae and 6-mo adult fish was automatically recorded and analyzed. For each recording, a total of nine motor parameters were considered. The statistics for the individual parameters are reported in Figure S10. Because a complex set of changes may go unnoticed when considered in isolation, logistic regression was used to study the dependence of the genotype, a categorical variable with only two possible outcomes (“mzLrrk2” or “wt”), from a combination of the motor parameters. This is the same as using overall swimming performance as indicator of loss of *lrrk2*. The results are summarized in Table 2; each model’s goodness of fit is provided in Figure S11.

**Table 2.** Multiple logistic regression analysis of the association of the swimming performance with the loss of *lrrk2*.

| time point | independent variable | estimate ($\beta$) | OR ($e^\beta$) | 95% CI | P |
| --- | --- | --- | --- | --- | --- |
| 5 dpf | inactive phase distance | −0.841 | 0.432 | 0.258–0.721 | 0.0014 |
|  | normal phase entry count | 0.216 | 1.242 | 1.071–1.439 | 0.0041 |
|  | normal phase duration | −0.050 | 0.951 | 0.916–0.988 | 0.0091 |
|  | bursting phase duration | −0.269 | 0.764 | 0.628–0.929 | 0.0071 |
| 10 dpf | inactive phase entry count | −0.071 | 0.932 | 0.873–0.994 | 0.0316 |
|  | inactive phase duration | −0.033 | 0.968 | 0.948–0.988 | 0.0018 |
|  | inactive phase distance | 0.521 | 1.684 | 1.149–2.468 | 0.0076 |
|  | normal phase duration | 0.064 | 1.066 | 1.014–1.121 | 0.0126 |
|  | normal phase distance | −0.151 | 0.860 | 0.792–0.933 | 0.0003 |
|  | bursting phase duration | −0.166 | 0.847 | 0.735–1.976 | 0.0219 |
|  | bursting phase distance | 0.177 | 1.194 | 1.079–1.321 | 0.0006 |
| 6 mo | normal phase distance | 0.008 | 1.008 | 1.004–1.011 | <0.0001 |
|  | bursting phase distance | 0.003 | 1.003 | 1.002–1.004 | <0.0001 |

The original motor variables (inactive/normal/bursting phase entry count/duration/distance for 5- and 10-dpf larvae; normal/bursting phase entry count/duration/distance for 6-mo adults) were subjected to backward stepwise elimination to identify the best fitting model at each time point. The selected variables and relative statistics are shown. For each variable, the odds ratio (OR) represents the change in the relative probability ( $p$ ) of a recorded animal to be *mzLrrk2* ( $p=1$ ) over *wt* ( $p=0$ ) for an $x$ -unit change, the other variables held constant. Unit changes of $x=10$ for phase entry count and phase distance, $x=1$ for all other variables were deemed opportune. The estimated 95% confidence interval for each OR is reported. The intercept, i.e. the estimated OR when all covariates equal 0, is omitted.

At 5 dpf, the estimated odds ratios revealed that larvae were more likely to be mzLrrk2 if swimming more frequently at the normal speed range (normal phase entry count, OR=1.242) while spending less time in bursting swim mode (bursting phase duration, OR=0.764). At 10 dpf, mzLrrk2 larvae tended to alternate inactive to bursting phases (inactive phase distance, OR=1.684; bursting phase distance, OR=1.194), rather than swimming at the normal speed range (normal phase distance, OR=0.860), with lower cumulative bursting phase duration (OR=0.847). This “stop-and-go” swimming pattern might reflect an impaired ability to initiate or sustain movements, similar to bradykinesia in PD patients. However, at 6 mo, although the distance swum in either normal or bursting mode was a very strong predictor (*P*<0.0001 for both), the estimated odds ratios were very close to 1 (OR=1.008 and OR=1.003, respectively), indicating that the difference between adult mzLrrk2 fish and controls was very subtle.

Several non-motor symptoms often precede motor disease and aggravate disability in later stages of PD pathology. Anxiety and olfactory dysfunction are amongst the most prevalent in PD patients both after and prior to diagnosis [56]. Therefore, anxiety levels and response to olfactory stimuli were analyzed in fish. To test anxiety levels, two paradigms were used: (i) wall-hugging behavior, or thigmotaxis, for both larvae and adults (Figure S12a–d), and (ii) dark-to-light preference, or scototaxis, for adults only (Figure S12e). To evaluate the overall olfactory function in adult fish, the response to an amino acid mixture odorant stimulus was measured (Figure S12f). None of the assays revealed any significant difference between mzLrrk2 fish and controls.

## Discussion

Here we present the characterization of the brain phenotype of a complete zebrafish *lrrk2* knockout model (Figure 7). The homology between human LRRK2 and zebrafish Lrrk2 proteins (Figure 1a), the temporal and regional dynamics, and entity of *lrrk2* gene expression in the zebrafish brain from early embryo to adult (Figure S1, 2) make the zebrafish a suitable tool to study LRRK2 biology. The complete deletion of the entire *lrrk2* locus (Figure 1b) furthermore provides a genetically unambiguous tool to study LRRK2 deficiency in a vertebrate *in vivo*. The importance of this feat is twofold. First, because human LRRK2 variants are often considered to be gain-of-functions, great effort is being invested in the generation and evaluation of LRRK2 inhibitors [57]. However, pathogenicity of *Lrrk2* knockout in peripheral organs in rodents raise the possibility of alternative or additional loss-of-function contributions to disease pathology, as exemplified also by our examination of the *Lrrk2*-null phenotypes. In addition, a *lrrk2*-null zebrafish model is potentially applicable for high-throughput drug screening. In the context of PD modeling, alternatives to the gain-of-function hypothesis have been suggested before. Although human *LRRK2* nonsense mutations have been reported, though never associated with disease [38, 58], there is evidence supporting hypomorphic [25], dominant-negative [21, 23], and even protective [18, 59] effects of established PD-associated LRRK2 mutations. In fact, it has been hypothesized that both “too much” or “too little” LRRK2 might detrimentally converge on synaptic function [60]. Being a completely amorphic model, the mzLrrk2 zebrafish model reported here represents a genetically unambiguous situation that provides novel opportunities for current and future investigations on LRRK2 loss-of-function.

**Figure 7.**
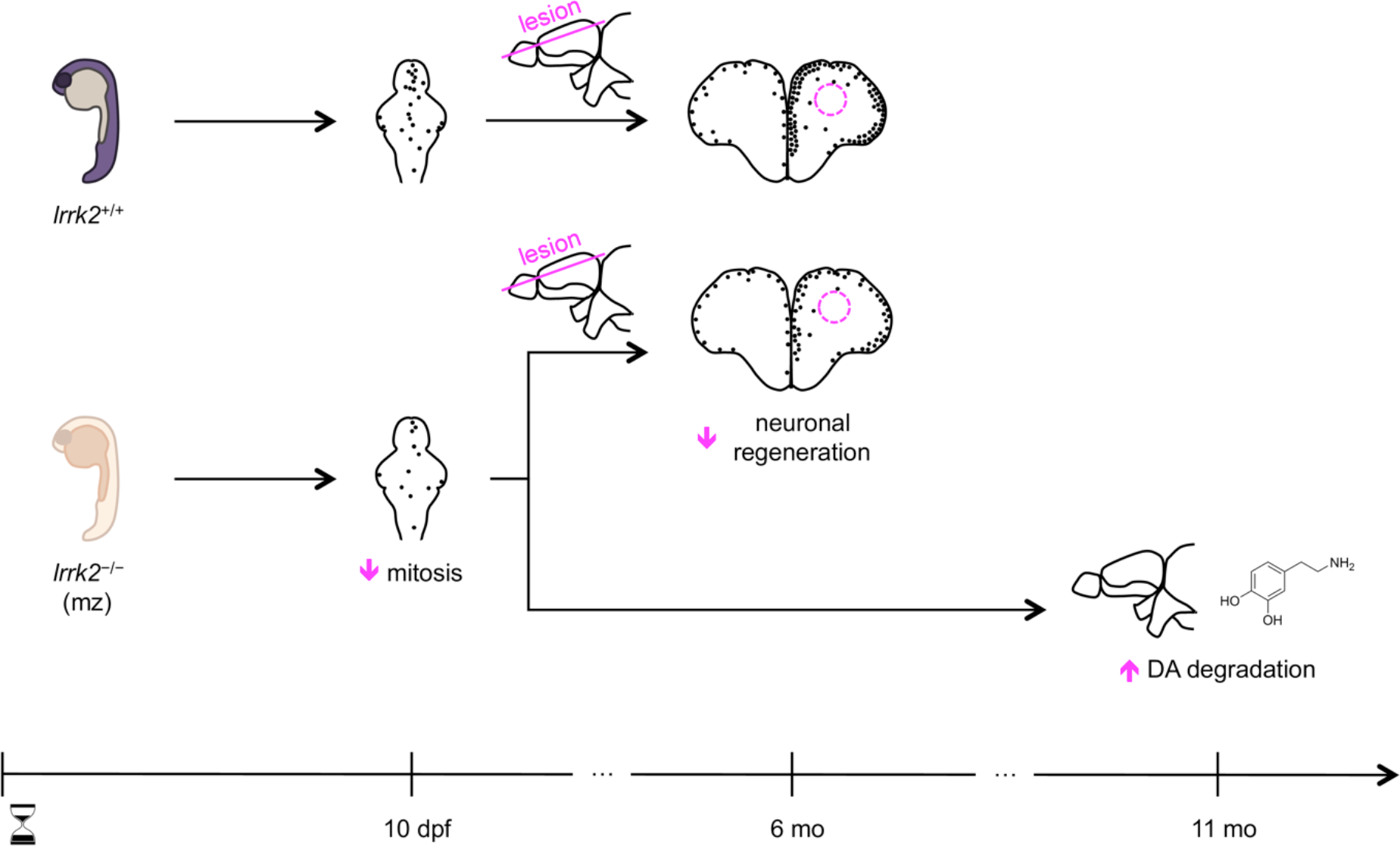
Overview of the main findings.

In contrast with published MO-induced phenotypes [32, 34], we show that zebrafish *lrrk2* is a resilient locus (Figure 2–4). However, in contrast with previous Lrrk2 knockout studies in mice [29], we show that loss of *lrrk2* does have specific effects on the brain. In particular, mzLrrk2 larvae initially display reduced CA neurons and a combination of distinct motor signatures resembling hypokinesia, that however resolve spontaneously during further development. A possible reason for the lack of an overt PD-like phenotype in mzLrrk2 fish may be the lack of a zebrafish α-synuclein ortholog. Although α-synucleinopathy is not pathognomonic of PD, not even in *LRRK2* mutation carriers, progression of Lewy body pathology correlates with non-motor symptoms [3], including psychiatric disturbances and olfactory dysfunctions [61]. On these grounds, the normal anxiety levels and olfactory responsiveness in mzLrrk2 zebrafish that we observed seem reasonable (Figure S12). Yet, no PD-like brain pathology manifests in either transgenic [7-12] or knockout [29] *LRRK2* mouse models, where α-synuclein is present.

Strikingly, we observe perturbed amine catabolism, most prominently increased dopamine degradation, in 11-mo mzLrrk2 fish (Figure 4k). This result is particularly meaningful considering that brain dopamine and/or catabolite levels are normal in mice lacking endogenous [28, 29] or overexpressing human wild-type LRRK2 [11], but are reduced in mice expressing pathogenic LRRK2 variants, although the effect may be regional [10, 11] or transgene-dependent [11]. The neurochemical signatures in these mouse models have been difficult to interpret, because they are not clearly linked to problems in either dopamine synthesis [10] or turnover [10, 11]. In contrast, we specifically find enhanced monoamine oxidase (MAO)-dependent amine degradation in mzLrrk2 fish. How this effect relates to loss of *lrrk2* is currently unclear, and hence it might be indirect. Previous observations have shown that both expression of the pathogenic LRRK2 R1441C variant [12] and *LRRK2* knockdown [62] impair synaptic transmission. If this is the case also for mzLrrk2 fish, increased MAO activity may be a protective mechanism against dopamine accumulation either at the presynaptic terminal, due to impaired neurotransmitter release, or at the synaptic cleft, due to impaired reuptake, as excess dopamine represents an oxidative threat due to radical formation [63]. However, while removing excess dopamine, MAO also generates toxic hydrogen peroxide [63], being potentially deleterious in the long term. Given the clinical relevance of MAO inhibitors as a treatment for PD [64], our findings highlight the importance to clarify the role of MAO given a predisposing genetic background.

An interesting question is whether mzLrrk2 fish may be protected, to some extent, from neurodegeneration by their potential to regenerate the brain after lesion [54, 55]. We provide compelling evidence that loss of *lrrk2* reduces the mitosis rate in the brain. The phenotype is specific to maternal-zygotic animals. Accordingly, *ad hoc* formation of cells to replace lost neurons is unlikely. However, this does not rule out that pro-regenerative cues may be present. Conceivably, the suppression of pro-regenerative mechanisms might aggravate the phenotype. One such mechanism may be protective autoimmunity [65]. Treatment with dexamethasone, a steroid anti-inflammatory drug, has been shown to dampen the regenerative ability of the zebrafish central nervous system [66, 67]. We verified that chronic exposure to dexamethasone has no influence on cell proliferation in 10-dpf mzLrrk2 brains (Figure S13). Therefore, if a regenerative response were underway, this would be independent of neuroinflammation. Other forms of plasticity might however come into play, including transdifferentiation [68]. Consequently, longitudinal studies will be needed to search for cell loss in aging fish.

Finally, we suggest that perturbing assays, such as the brain lesion we employed here, may reveal aspects of Lrrk2 pathobiology that are otherwise unnoticeable in unchallenged mzLrrk2 fish. In particular, we exploited the zebrafish as a regenerating organism and tested the brain reparative potential upon stabbing the adult telencephalon [54], and indeed, we observe impaired neuronal regeneration (Figure 6c–c’’). We cannot rule out whether this conditional phenotype is an indirect consequence of the hypoproliferative phenotype we observe in the larval brain. Alternatively, the role of Lrrk2 in the control of cell proliferation may be more general during development, and more specialized later on in adult brains. Such a changing role could be reflected by *lrrk2* expression in the brain, which switches from being ubiquitous during development (Figure S1) to being more confined to specific areas in the adult brain, notably including the telencephalic neurogenic niche (Figure S2b–d). In either case, priming or supporting mechanisms might be required for Lrrk2 to partake in neuronal regeneration. Neuroinflammation appears as a promising candidate [44]. Of note, cultured rat microglia exhibit increased *LRRK2* expression upon acute inflammation and impaired inflammatory response upon *LRRK2* knockdown [69]. Furthermore, microglia in BAC human LRRK2 G2019S transgenic mice subjected to stab-wound or laser-injury display impaired ability to isolate the lesion site [70]. The zebrafish Lrrk2 may have a similar pro-inflammatory role, as suggested by our finding of reduced leukocytosis in larvae after TPA-induced systemic acute inflammation (Figure 3d, d’). Because acute inflammation is required for neuronal regeneration [66], a plausible scenario would then be that impaired production of neural precursor cells combined with defective microglia in mzLrrk2 fish impede brain repair. Transferred to humans, impaired LRRK2 function may lead to neurodegeneration through the accumulation of unresolved damages such as the ones following traumatic brain injury. Albeit recognized alone as a risk factor for PD [71], traumatic brain injuries are yet to be examined in conjunction with known genetic factors, including *LRRK2* mutations. Targeted analyses on animal models and epidemiological studies are therefore warranted. To summarize, our results show that loss of *lrrk2* can compromise specific vertebrate brain functions, including MAO-dependent amine catabolism and regenerative capacity upon lesion. We believe that similar defects in humans might play a contributing role in the prodromal stages of PD and are worth investigating further. To this aim, our zebrafish *lrrk2* knockout model offers a unique possibility to study *in vivo* the consequences of LRRK2 loss of function in a regenerating vertebrate system and, coupled with the well-established high-throughput screening amenability of zebrafish, provides a means for identifying targets of interest in a fish-to-mammal translational perspective.

## Methods

### Ethics statement

All animal experiments were conducted according to the guidelines and under supervision of the Regierungspräsidium Dresden (permits AZ: 24D-9168.11-1/2008-2; AZ: 24D-9168.11-1/2008-3; AZ: 24-9168.11-1/2013-5; AZ: 24-9168.11-1/2013-14; DD24-5131/346/11; DD24-5131/346/12). All efforts were made to minimize animal suffering and the number of animals used.

### Sequence alignment analyses

Sequences were retrieved from the latest assemblies of the human and zebrafish genomes (GRCh38p.7 and GRCz10, respectively). Alignment analyses were performed using Clustal Omega [72] and BLAST [73].

### Cas9 and gRNA construction

*Cas9* mRNA and gRNAs were synthesized as previously described [74]. Briefly, *Cas9* mRNA was synthesized by *in vitro* transcription using T3 mMESSAGE mMACHINE kit (Ambion). gRNAs were generated and purified using the MEGAshortscript T7 and *mir*Vana miRNA isolation kits (Ambion), respectively. Sequences of the genomic target sites and oligonucleotides for making gRNAs are listed in Table 3.

**Table 3.**
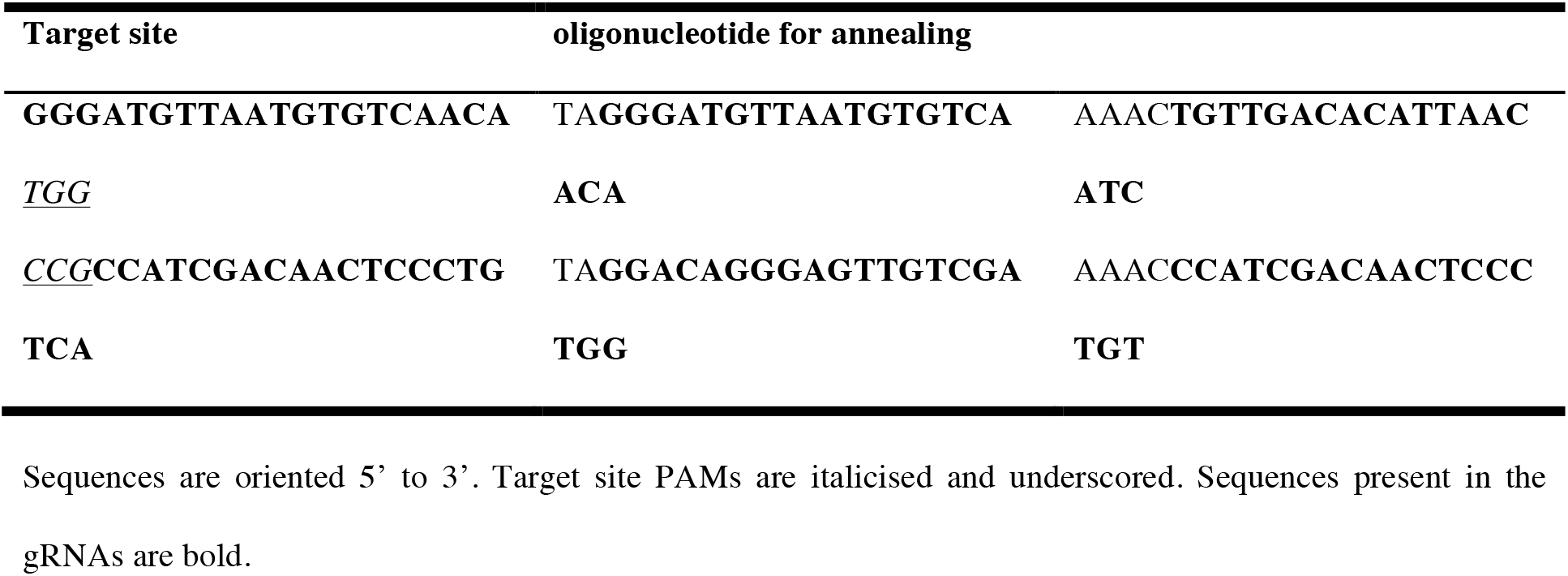
gRNA target sites for *lrrk2* locus deletion.

### Plasmid construction

A *lrrk2* cDNA fragment (c.3009_7130 according to the Human Genome Variation Society guidelines [39]) flanked by BamHI (5’) and ClaI (3’) restriction sites was cloned into a *pCS2+MT* vector [75]. The *lrrk2* rescue fragment including the 5x*Myc* repeat, the TAG stop codon, the poly(A) signal, and flanked by NheI (5’) and AscI (3’) restriction sites was cloned into the vector *pTol(hsp70l:mCherry-T2A-CreER^T2^
*) [76] to result in the simultaneous expression of the corresponding Lrrk2 protein fragment and the mCherry reporter under the control of the heat-inducible *hsp70l* promoter. Primers used for cloning are reported in Table 4.

**Table 4.**
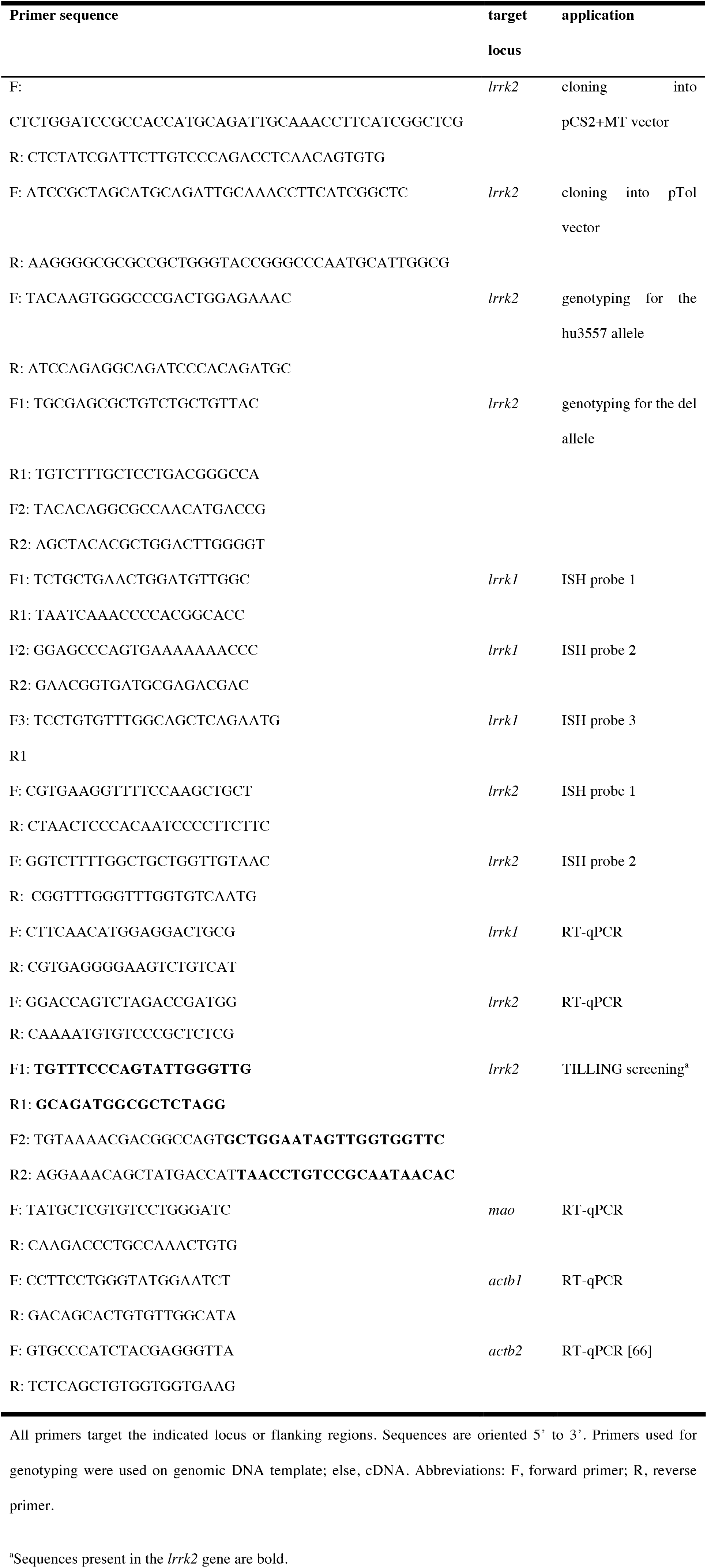
Primers used.

### TILLING and TALEN screening

Targeting induced local lesion in genomes (TILLING) mutagenesis screening was performed as previously described [53]. Outer (F1, R1) and inner (F2, R2) primers used are reported in Table 4. During the screening the allele *hu3557*, a splice mutation affecting the splice donor of *lrrk2* exon 27 (c.3972+2T>C) was recovered and designated as “tud112”. We also generated an additional, early stop codon-carrying *lrrk2* allele (c.1980_1990del) using transcription activator-like effector nucleases (TALENs), denominated *tud115*, which however could give rise to a truncated Lrrk2 product from a downstream alternative translation initiation site, and was therefore not considered further.

### Zebrafish husbandry and germ line transformation

Zebrafish were raised and maintained as previously described [77]. Zebrafish embryos were obtained by natural spawnings of adult fish and staged according to hours post fertilization (hpf) or standard criteria [78]. The wild-type line used was AB. For CRISPR/Cas9-mediated mutagenesis, 150 ng/µL dual NLS-tagged zebrafish codon-optimized *cas9* mRNA/50 ng/*µ*L gRNAs/0.2% phenol red were co-injected into fertilized eggs, the embryos raised to adulthood, crossed to AB wild-type fish and the resulting F1 embryos screened by PCR. The identified mutation (c.−61_*42del; see Results section) was designated as “tud113”. For germ line transformation, plasmid DNA (*pTol(hsp70l:mCherry-T2A-lrrk2(c.3009_7130)-Myc)*) and transposase mRNA were injected into fertilized eggs (F0), raised to adulthood and crossed to AB wild-type fish as previously described [79]. To identify transgenic carriers, undechorionated F1 embryos at 20 hpf were heat-shocked, examined under a fluorescent microscope after a 4 h waiting period and positive embryos were raised. The established line was designated as “tud114”.

### Genotyping

Genotyping was performed using genomic DNA from individual or pooled embryos/larvae or fin clips from larvae or adult fish. Primers used are listed in Table 4. For genotyping for the *lrrk2^tud113^
* allele, see Results section. For genotyping for the *lrrk2^tud112^
* allele, see Figure S7.

### Animal experiments

Fin clipping for genotyping purposes was performed on adult fish following anesthesia with 0.02% tricaine. Drug treatments are summarized in Table 5. Heat shocks were administered daily from 1 to 10 days post-fertilization (dpf). Treatment consisted in sudden exposure to 42 ºC E3 medium, followed by 4 h incubation at 37 ºC. Fish were sacrificed 4 h after the last heat shock. Stab lesions were performed on male adult fish as previously described [54].

**Table 5.** Summary of drug treatments.

| **Drug** | **dose and solvent** | **exposure** |
| --- | --- | --- |
| BrdU | for larvae: 10 μM, E3 medium | 3 h |
| | for adults: 5 μM, fish water | 24 h |
| Dexamethasone | 10 μM from 25 mg/mL MetOH stock, autoclaved E3 medium | 9 d<sup>a</sup> |
|  | control: MetOH equivalent, autoclaved E3 medium |  |
| TPA | 10 ng/ml from 50 μg/mL DMSO stock, E3 medium; | 2 h |
|  | control: DMSO equivalent, E3 medium |  |

All drugs were purchased from Sigma. Abbreviations: BrdU, 5-bromo-2'-deoxyuridine; DMSO, dimethyl sulfoxide; MetOH, methanol; TPA, 12-*O*-tetradecanoylphorbol-13-acetate.
<sup>a</sup>From 1 to 10 dpf, solution changed daily.

### Behavioral analyses

The ZebraBox and ZebraCube apparatus were used in combination with the Viewpoint Application Manager software. All recordings were performed on individually isolated animals between 2–6 pm. Larvae at 4 dpf were transferred into 24-well plates and therein grown until 10 dpf. Each well was internally lined with Parafilm to minimize reflection and filled with 750 µL E3 medium, changed daily. Animals whose tracking was lost by the software for over 20% of the total recording time were excluded from the analyses. To quantify spontaneous swimming and thigmotaxis, adult fish were lodged in opaque cylindrical boxes (ø=80 mm) filled with 100 mL fish facility water, else in opaque parallelepipedal boxes (l×w=190×80 mm) filled with 500 mL fish facility water. Each recording was preceded by 10 min acclimatization inside the apparatus. Spontaneous swimming was assessed for 10 min (integration period: 600 s) in the dark under infrared light. Appropriate speed thresholds were chosen based on developmental stage: 2–10 mm/s for larvae; 2–40 mm/s for adults. Based on the speed thresholds, three swimming phases were defined: inactive phase, below the lower threshold; normal swimming phase, between the lower and upper threshold; bursting phase, above the upper threshold. For each swimming phase, three parameters were considered: entry count, duration (s), and distance swum (mm). Thigmotaxis was assessed using the same recordings of spontaneous swimming activity. To this aim, the recording arena was digitally subdivided into an outer and inner area (for larvae ø=15.6/10.6 mm; for adults ø=80/55 mm). Scototaxis was assessed for 10 min in half-black, half-white parallelepipedal boxes. Olfactory function was assessed by delivering a stimulus in either of the shorter sides of parallelepipedal boxes. The stimulus consisted in 0.6 mL of an amino acid mix (Ala, Cys, His, Lys, Met, Val, 0.1 mM each) delivered through a syringe pump (1.5 mL/min). Fish were starved for 24 h before the experiments. Fish behavior was recorded 5 min before and 5 min after stimulus delivery. For every 1 min of recording, a preference index was defined [80] as
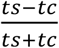
, where ts is the time spent in the stimulus side, tc the time spent in the control side.

### Tissue processing and histochemistry

Tissue processing is summarized in Table 6. Bleaching was performed using 3% hydrogen peroxide/0.1% Tween20/1% potassium hydroxide. Clearing [81] was carried out overnight at room temperature. The procedure caused slight swelling of tissue, more pronounced in younger samples. Immunohistochemistry (IHC) was performed as previously described on whole embryos/larvae [82], dissected larval brains [46], and cryosections [83]. Primary antibodies used are listed in Table 7. Secondary antibodies (1:500) were conjugated to Alexa Fluor 488, 555, 633, 700 (Invitrogen). For combined staining with one or more antibodies requiring antigen retrieval the following precautions were used: HuC/D IHC was performed first if combined with BrdU IHC, otherwise last; PCNA, TH(1) IHC were performed last. TUNEL assay on larval brains was carried out using the ApopTag Red or Fluorescein *In Situ* Apoptosis Detection Kits (Millipore) according to the manufacturer’s instructions with the following adjustments: tissue was washed 3×10 min with sodium citrate/Triton X-100 0.1% in PBS prior to acetic acid/ethanol post-fixation; incubation in the equilibration buffer was carried out for 1 h at room temperature. When combined with IHC, TUNEL assay was performed first. *In situ* hybridization (ISH) and probe generation were performed on embryos and at least three adult individuals [84]. All synthesized and individually tested probes (Table 4) showed a redundant pattern. Monoamine oxidase histochemistry (MAO HC) was carried out as previously described [46] with the following adjustments: staining was developed for 90 min; stained tissue was post-fixed in 4% paraformaldehyde.

**Table 6.**
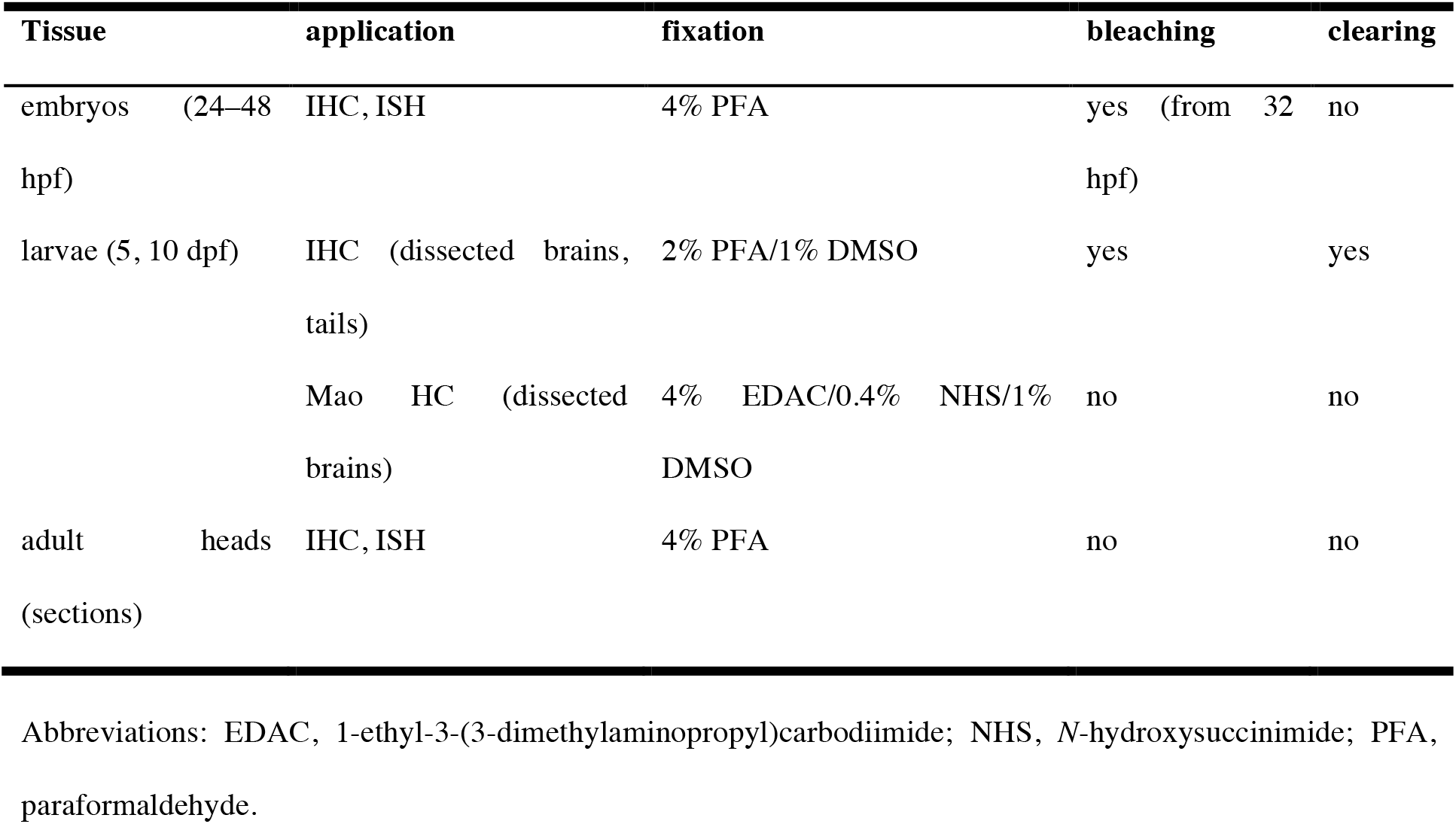
Summary of tissue processing.

**Table 7.**
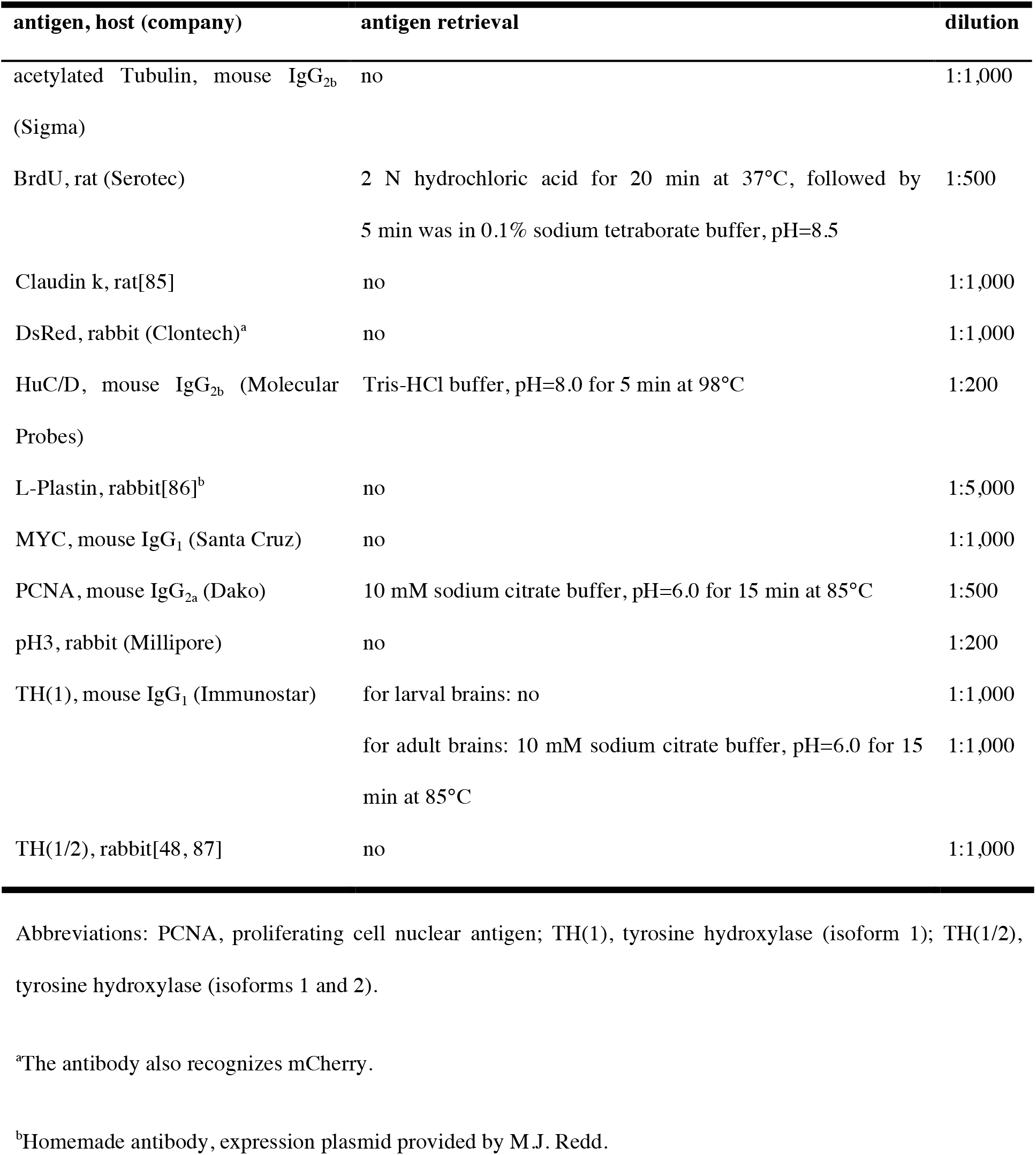
Primary antibodies used.

### High performance liquid chromatography measurements

Tissue samples consisted each in 10-pooled whole larvae or single adult brain. Larvae were starved 24 h before tissue collection to minimize possible contamination from amines in the gastrointestinal tract. An equal number of male and female adult fish were sacrificed. Tissue homogenization and catabolite measurements via electrochemical detection coupled with high performance liquid chromatography (HPLC) were performed as previously described [46].

### Immunoblotting

Protein pellets obtained from the tissue samples used in HPLC were homogenized by sonication at room temperature in 5% SDS in PBS, pH 7.4, to perform the BCA assay. The solutions were then diluted with a loading buffer stock containing Tris-HCl, pH 6.8, β-mercaptoethanol, glycerol, and bromophenol blue (final concentration: 0.0625 M Tris -HCl, 5% β-mercaptoethanol, 10% glycerol, and 0.002% bromophenol blue). The samples were heated at 95°C for 5 min and loaded on a gel for SDS-PAGE (4% stacking gel, 9% separating gel). The loading volume was adjusted in order to load 20 *µ*g total protein per well. PageRuler Prestained protein markers (Thermo Scientific) were used to control protein separation. The proteins were electrophoretically transferred to an Immobilon P PVDF membrane in an eBlot device (GenScript) according to the manufacturer’s instructions and the membrane was processed for immunoblotting in an Odyssey CLx system (Li-Cor) according to the manufacturer’s instructions. Mouse monoclonal anti-TH(1) antibody (Immunostar) diluted was used as the primary antibody (1:2,000), and a goat anti-mouse IRDye800 antibody (Li-Cor) was used as the secondary antibody (1:20,000). The fluorescent bands were quantitated by the ImageStudio software supplied with the Odyssey CLx system, and the membranes were stained with ProAct membrane stain (M282–1L, Amresco Inc.) as the loading control.

### MAO activity assay

Peroxidase-linked colorimetric assay of MAO activity was performed as previously described [88].

### RNA extraction and RT-qPCR

Total RNA was isolated from pooled embryos using TRIzol (Invitrogen). cDNA was generated using Superscript III First Strand Synthesis System (Invitrogen). Reverse transcriptase-quantitative PCR (RT-qPCR) was performed using the LightCycler 480 SYBR Green I Master mix and the LightCycler 480 Instrument (Roche). Target gene expression was normalized by *actb1* or *actb2* expression.

### Image acquisition and processing

Confocal images were acquired with a Zeiss LSM 780 upright confocal microscope using C-Apochromat 10x/0.45 W and LD LCI Plan-Apochromat 25x/0.8 Imm Corr DIC M27 objectives for water immersion. Bright-field images were acquired with an Olympus DP71 or DP80 color cameras connected to an MVX10 microscope. All devices were provided by the BIOTEC/CRTD Light Microscopy Facility. Images were processed using Fiji [89]. Processing was applied equally across entire images and to controls. Cell quantification was carried out manually through whole stacks. To analyze microglia/leukocyte morphology and complexity, confocal stacks were background-subtracted using the sliding paraboloid method, despeckled, and thresholded using Li’s method (Figure S4a–a’’’). Obvious artifacts were manually removed from subsequent processing and analyses. The 3D ImageJ Suite plugin [90] was used to segment objects (minimum size threshold: 1,000; objects on borders excluded) and extract morphological data from 3D masks. The same 3D masks were subsequently skeletonized and subjected to 3D skeleton analysis using the AnalyzeSkeleton plugin on Fiji [91]. Loops were pruned using the shortest branch method. For each skeleton, the longest shortest path was also calculated. A ramification index was defined as 
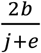
 where b is the number of branches, j the number of junctions, e the number of end-points, as previously defined [91].

### Statistical analyses

The data analysis for this paper was generated using: the Real Statistics Resource Pack software (Release for Mac 3.1.2, copyright 2013–2016) developed by C. Zaiontz (www.real-statistics.com); R [92]; and GraphPad Prism version 7.0b for Mac OS X. To compare means, requirements of normal distribution and homoscedasticity were checked using Shapiro-Wilk’s test and the F-test, for two groups, or Levene’s test, for more than two groups, respectively. To determine the statistical significance of group differences, *P* values were calculated using: Student’s *t*-test or ANOVA, for normally distributed and homoscedastic data; Student’s t-test with Welch’s correction, for normally distributed and heteroscedastic data; Mann-Whitney’s *U*-test for non-normally distributed data. Multiple comparisons following ANOVA were performed using Dunn-Šidák’s method. For multivariate logistic regression analyses, the best fitting models were automatically selected via backward stepwise elimination. The model performance was visualized and assessed using the methods previously described [93]. For each experiment, sample sizes are reported in the Figures. Plot features are described in the Figure legends. Within the Figures, significant comparisons are marked by asterisks: *, *P*<0.05; **, *P*<0.01; ***, *P*<0.001; ****, *P*<0.0001. *P* values rounded to three decimal places are reported in the Figure graphs for values comprised between 0.050 and 0.059; *P* values rounded to four decimal places are reported in the main text.

## Acknowledgments

The authors express thanks to: our histology facility, light microscopy facility, and fish facility for excellent support; M. Reimer (CRTD) for providing the anti-Claudin k antibody; M.J. Redd (Department of Pathology, University of Utah, Salt Lake City, Utah, USA) for providing the anti-L-Plastin antibody; J. Guck (BIOTEC) for providing the syringe pump; J. Sterneckert, G. Kempermann, and W. Schlechte-Wełnicz (CRTD) for comments on the manuscript, and our funding agencies, as indicated, for support.

